# Crash landing of *Vibrio cholerae* by MSHA pili-assisted braking and anchoring in a viscoelastic environment

**DOI:** 10.1101/2020.07.05.188680

**Authors:** Wenchao Zhang, Mei Luo, Chunying Feng, Huaqing Liu, Hong Zhang, Rachel R. Bennett, Andrew S. Utada, Zhi Liu, Kun Zhao

## Abstract

Mannose-sensitive hemagglutinin (MSHA) pili and flagellum are critical for the surface attachment of *Vibrio cholerae*, the first step of *V. cholerae* colonization on host surfaces. However, the cell landing mechanism remains largely unknown, particularly in viscoelastic environments such as the mucus layers of intestines. Here, combining the cysteine-substitution-based labelling method with single-cell tracking techniques, we quantitatively characterized the landing of *V. cholerae* by directly observing both pili and flagellum of cells in a viscoelastic non-Newtonian solution consisting of 2% Luria-Bertani and 1% methylcellulose (LB+MC). The results show that MSHA pili are evenly distributed along the cell length and can stick to surfaces at any point along the filament. With such properties, MSHA pili are observed to act as a brake and anchor during cell landing which include three phases: running, lingering, and attaching. Importantly, loss of MSHA pili results in a more dramatic increase in mean path length in LB+MC than in 2% LB only or in 20% Ficoll solutions, indicating that the role of MSHA pili during cell landing is more apparent in viscoelastic non-Newtonian fluids than viscous Newtonian ones. Our work provides a detailed picture of the landing dynamics of *V. cholerae* under viscoelastic conditions, which can provide insights into ways to better control *V. cholerae* infections in real mucus-like environment.

## Introduction

*Vibrio cholerae*, a human pathogen that causes the debilitating disease cholera, is a natural inhabitant of aquatic ecosystems (*Almagromoreno et al., 2015; Kaper J B, 1995*). They can form biofilms on both biotic and abiotic surfaces, which increases their infectivity and environmental survival (*Donlan et al., 2002; Silva et al., 2016; Teschler et al., 2015; Yildiz et al., 2009*).

Bacterial appendages have been shown to play important roles in regulating bacterial activities especially biofilm formation during microbe-host interactions. The flagellum is required for biofilm formation in a variety of bacteria species, such as *E. coli* (*Pratt et al., 1998*), *P. aeruginosa* (*O’Toole et al., 1998*), and *V. cholerae* (*Guttenplan et al., 2013; Watnick et al., 1999*). Mutants lacking flagella in both *E. coli* and *Vibrio vulnificus* have been observed to be defective for attachment (*Friedlander et al., 2013; Lee et al., 2004*). Type IV pili (TFP) are another type of filamentous appendages commonly found on many bacteria and archaea, which have diverse functions such as cellular twitching motility, biofilm formation, horizontal gene transfer, and host colonization (*Piepenbrink et al., 2016*). *P. aeruginosa* display two types of TFP-driven twitching motility (*Gibiansky et al., 2010*). *Neisseria gonorrhoeae* have shown a TFP-dependent attachment, leading to the formation of microcolonies on host cell surfaces (*Higashi et al., 2007*). In contrast, although *V. cholerae* biosynthesize three types of TFP that are expressed under different scenarios, they have not been observed to twitch on surfaces. These three pili are: chitin-regulated competence pili (ChiRP; formerly termed PilA), toxin co-regulated pili (TCP), and mannose-sensitive hemagglutinin type IV pili (MSHA) (*Meibom et al., 2004; Reguera et al., 2005; Yildiz et al., 2009*). ChiRP pili were observed to be able to grasp extracellular DNA and transport it back to the cell surface via pili retraction (*Ellison et al., 2018*). TCP pili are important for host colonization and pathogenesis (*Kirn et al., 2000; Thelin et al., 1996*). In contrast to these two types, MSHA pili are known to be important for surface attachment of *V. cholerae* (*Utada et al., 2014; Watnick et al., 1999*).

Motility has been shown to be a crucial element for *V. cholerae* colonization of the epithelium, leading to successful infection of the human host (*Krukonis et al., 2003; Tsou et al., 2008*). Two types of near-surface motility, roaming and orbiting, were observed in *V. cholera* (*Utada et al., 2014*). It has been further suggested that *V. cholerae* synergistically employ the use of their flagella and MSHA pili to enable a hybrid surface motility that facilitates surface selection and attachment (*Utada et al., 2014*). However, there is a lack of direct observational evidence of the appendages in question. More importantly, the environmental niches *V. cholerae* encounter in their life cycle typically include viscoelastic mucus (*Almagromoreno et al., 2015*). The mucus layer of animal intestines is estimated to have a wide range of viscosities, varying anywhere from the viscosity of water (∼1 cP) to 1000-fold higher (1000 cP) (*Lai et al., 2009*). How cells land on surfaces in such viscoelastic environments is still not clear. To answer these questions, direct live-cell visualization of the pili and flagellum in real-time in viscoelastic conditions is needed.

Recently, there have been significant advances in techniques for directly observing cell appendages (*Blair et al., 2008; Ellison et al., 2019; Ellison et al., 2018; Ellison et al., 2017a; Nakane et al., 2017; Renault et al., 2017; Skerker et al., 2001; Talà et al., 2019*). Among them, the cysteine substitution-based labelling method is specific and has been successfully applied to visualize tight adherence (TAD) pili of *C. crescentus* and type IV pili of *V. cholera* (*Ellison et al., 2019*).

In this paper, by combining a cysteine substitution-based labelling method with single-cell tracking, we directly observed the individual pili and flagellum of landing cells in viscoelastic media and revealed the dynamic landing sequence of *V. cholerae* as it makes initial surface attachment. The role of MSHA pili during cell landing in viscoelastic environment is demonstrated. Our work provides a detailed picture of the landing dynamics of *V. cholerae* under viscoelastic conditions, during which, the synergistic functions of MSHA pili and flagellum are elucidated.

## Results

### MSHA pili are evenly distributed along cell length with a constant length density

To visualize the MSHA pili, we constructed a mutant (MshAT70C) by cysteine substitution, which can subsequently be labeled with highly-specific maleimide dyes (*Figure 1*a and *Figure 1-Figure supplement 1*), following the protocol in Ellison *et al*. (*Ellison et al., 2019; Ellison et al., 2017a*). To observe the distribution of MSHA pili on the cell surface, we simultaneously stained the plasma membrane with FM4-64 in Figure 1a.

**Figure 1.**
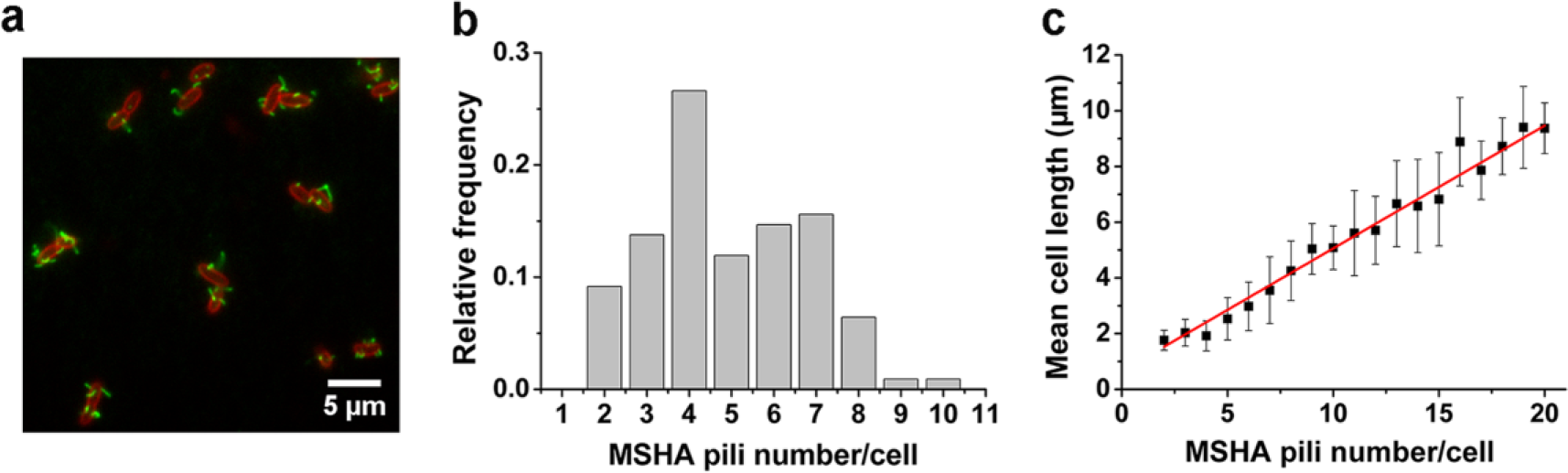
MSHA pili are evenly distributed along cell length with a constant length density. (a) Examples of labeled MSHA pili observed on cell bodies. Green fluorescence showing the AF488-mal labeled MSHA pili, red fluorescence showing the FM4-64 labeled plasma membrane. (b) Distribution of pili number per cell cultivated in LB medium. N_cell_ = 110. (c) The MSHA pili number per cell is linearly correlated with the cell length. Cells with longer length were obtained by 30∼50 min treatment using 10 μg/mL cephalexin. N_cell_ = 368. **Figure 1-source data 1. Figure 1b source data.** **Figure 1-source data 2. Figure 1c source data**. **Figure 1-Figure supplement 1. Labeling of *V. cholerae* MSHA pilus protein MshA with AF488-mal**. **Figure 1-Figure supplement 2. A 3D view of a typical *V. cholerae* cell showing the whole body distribution of MSHA pili; This cell has 6 pili**. **Figure 1-Figure supplement 3. MSHA pili labeling of the prepared samples**. To evaluate changes to the MSHA pilus during cell growth, the MSHA pili were labelled with two different colored dyes, AF546-mal (red) and AF488-mal (green), at 0 min and 40 min, respectively. (a) Representative double-color labeling image of MshAT70C cell, showing the new separate pilus (top left in green) and the secondary segments (lower left, green) at the end of the primary segments (lower left, red). Scale bar, 2 μm. (b) Representative double-color labeling image for a newly dividing cell, which is only labelled with AF488-mal. (c) In situ observation of MSHA pili growth stained at 0 and 50 min with AF488-mal. The results show that during a period of 50 min, the length of the cell changes from 3.0 μm to 5.1 μm, while the number of pili increases from 7 to 12. **Figure 1-Figure supplement 4. Hemagglutination assays**. MshAT70C point mutation does not affect MSHA pilus function. *V. cholerae* strains were grown in LB medium and assayed for MSHA production by hemagglutination. Two-fold dilutions of mid-log cultures of bacteria (left to right) were assayed for their ability to agglutinate sheep erythrocytes. Assay was repeated three times, and representative results are showed.

We visualized the positions of the different pili as the cell body rotates by recording high speed videos during surface landing (see one example of a three-dimensional model of a single cell reconstructed from the videos in *Figure 1-Figure supplement 2*). The results show evenly distributed MSHA pili along the cell length, indicating absence of preferred pili localization on the cell body. Quantitatively, we find that the majority of cells have approximately 3∼7 MSHA pili, with 4 MSHA pili per cell being observed most frequently, as shown in Figure 1b. These results are in agreement with recent reports (*Floyd et al., 2020*). Under our conditions, we observed MSHA pili growth (*Figure 1-Figure supplement* 3a and 3b) but no retraction.

The number of MSHA pili appears to be positively correlated with cell length since it increases as the cell grows (*Figure 1-Figure supplement* 3c). Statistically, the number of MSHA pili shows a linear relationship with cell length (Figure 1c), indicating that the length density of MSHA pili is roughly constant for *V. cholerae*.

### MSHA pili mediate *V. cholerae* landing by acting as a brake and anchor

The MSHA pili, which are uniformly distributed across the cell surface, play a crucial role in surface attachment of *V. cholerae* through pili-surface interactions (*Utada et al., 2014*). To elucidate the role of MSHA pili in the landing dynamics under viscoelastic conditions, we directly visualize the fluorescently labeled MSHA pili on *V. cholerae* swimming in a viscoelastic medium consisting of 2% LB and 1% MC (LB+MC).

Consistent with previous reports in normal aqueous solutions (*Utada et al., 2014*), the WT strain in LB+MC also exhibits orbiting behavior, characterized by multi-pass circular trajectories, and roaming behavior, characterized by highly exploratory, diffusive, trajectories. Typical roaming and orbiting trajectories in LB+MC are shown in Figure 2 (see more examples in *Figure 2-Figure supplement 1*). The roaming cell traces out a path that is linear trajectory over short distances, with a radius of gyration R_g_ =19.5 μm, and an average speed of 1.7 μm/s (see *Figure 2*a, b, Video 1). In contrast, the orbiting cell trajectory is much more circular with an average R_g_=1.6 μm and an average speed of 1.1 μm/s (see *Figure 2*c, d, Video 2). A 3D plot of speed plotted along the trajectory in both examples show that both phenotypes make momentary pauses, where their speed slows down; this can be seen clearly in Figure 2b, where the cell motion near a surface displays a characteristic alternation between moving and stopping (*Figure 2*b and 2d).

**Figure 2.**
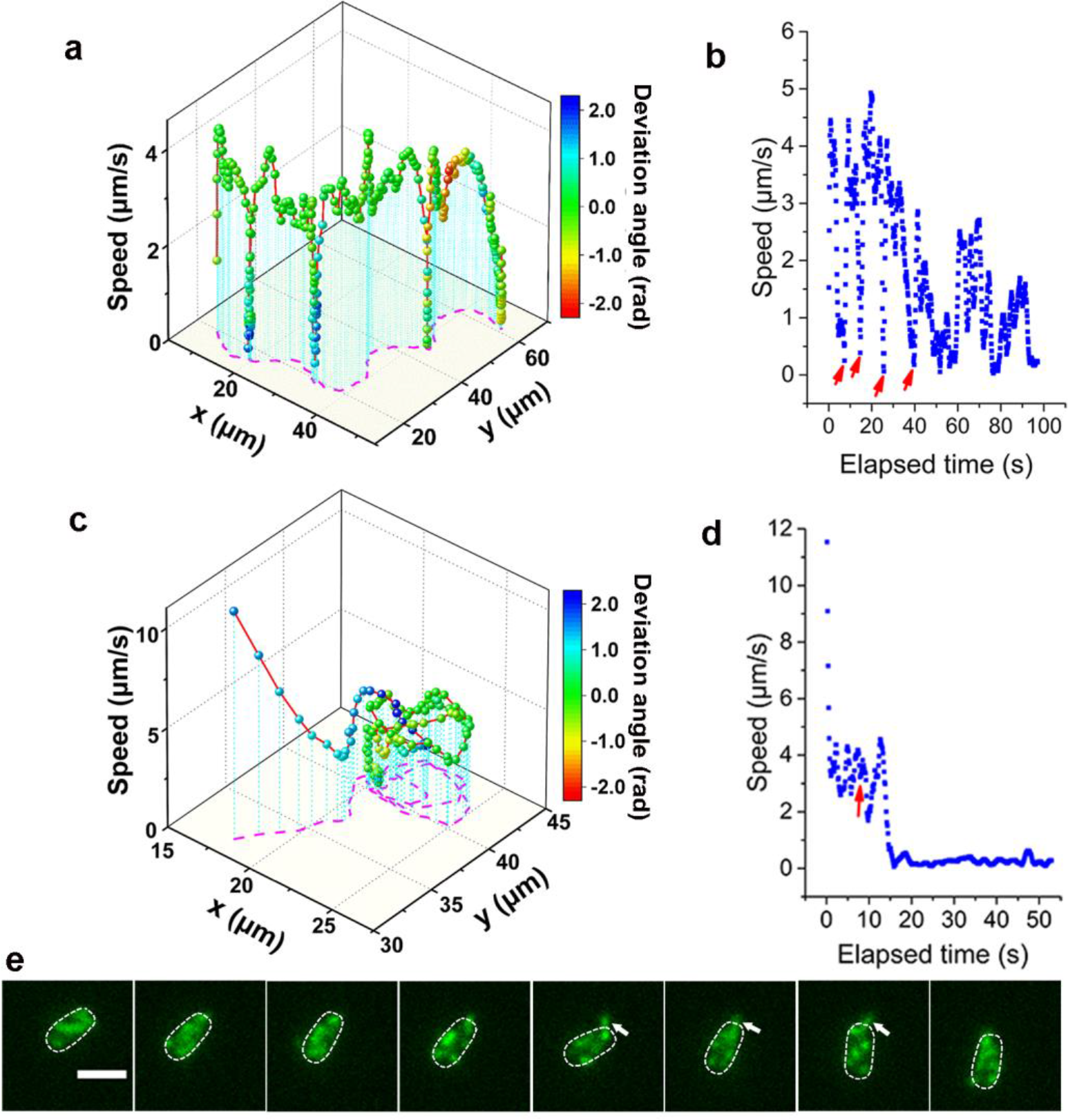
Analysis of roaming and orbiting, using cells of MSHA labelled MshAT70C. The 3D plot and speed changes over time of representative (a-b) roaming and (c-d) orbiting cells, respectively. The magenta dashed lines in panel (a) and (c) are the trajectories of cells and the color maps indicate the deviation angle. The arrows in panel (b) represent temporary attachment between MSHA pili and surface, where the speeds are close to 0. (e) Time-sequence snapshots of the orbiting cell in panel (c-d) at 130 ms intervals (see Video 2 for more details). The arrowheads show the stretched pilus, which corresponds to the red arrow in panel (d), indicating temporary attachment and stretching of pilus on the surface. Dashed lines indicate the estimated envelope of the cell body. Scale bar, 2 μm. **Figure 2-source data 1. Figure 2a source data**. **Figure 2-source data 2. Figure 2b source data**. **Figure 2-source data 3. Figure 2c source data**. **Figure 2-source data 4. Figure 2d source data**. **Figure 2-Figure supplement 1. Quantitative analysis of roaming and orbiting by MSHA labelled MshAT70C in 2% LB with 1% MC**. (a) Trajectories and (b) speed of typical roaming cells; (c) Trajectories and (d) speed of typical orbiting cells. **Figure 2-Figure supplement 2. Switch of temporary attached pili**. When transient pauses happened, the attached pilus could be switched from one to another or more. The arrows show the apparent pili attached with surface. Scale bar, 1 μm.

Such pauses are suggested to be caused by MSHA pili-surface interactions (*Utada et al., 2014*). However, by recording fluorescence video sequences, we directly visualized the process, thereby providing direct evidence that the pauses are due to transient contact between MSHA pili and surface. We show a transient pili-surface contact during orbiting in Figure 2e. In a sequence of frames, we see a transiently attached pilus become stretched due to cell motion away from the point of attachment. Subsequently, this pilus detaches from the surface as the cell continues to move, as indicated with the white arrowheads in Figure 2e (for more details, see Video 2). These results indicate that the MSHA pili can work as a brake to abruptly slow-down cell motion by transiently attaching to the surface. This is further confirmed by the observation that during the course of surface motion, different MSHA pili attach and detach, switching dynamically as the cell uses these as transient attachment points (*Figure 2-Figure supplement 2* and Video 3). Such a switching of the specific MSHA pili that are engaging the surface is caused by the rotation of cell body, which is required to balance the torque for flagellar rotation when cells swim. Thus, as the cell body rotates due to the rotation of the flagellar motor, different MSHA pili distributed on the cell body take turns approaching and receding from the surface. The switching of attached MSHA pili not only continues to slow-down cell motion but also changes the direction of motion. Taken together, this indicates that the pili distribution on the cell body may also affect cell-surface motion.

When the adhesion between MSHA pili and surface is sufficiently strong, the attachment point can act as an anchor point. We demonstrate this by showing the deflection of the trajectory of a swimming cell by the attachment of a, single, anchoring MSHA pilus; here, linear motion is bent into circular motion that is centered around the attachment point (see Video 4). We estimate the centripetal force for this motion to be on the order of 10^−21^ N, which is much smaller than the pN forces that pili can sustain (*Floyd et al., 2020; Maier et al., 2002*). The anchoring of MSHA pilus eventually leads to the irreversible attachment of the cell.

### The landing sequence of *V. cholerae* includes three phases

To further clarify the landing process, we labelled both flagellum and pili simultaneously using MshAT70CFlaAA106CS107C mutant. Figure 3 shows an example of the complete landing process of an orbiting cell. Based on the pattern of motion displayed by the cell (*Figure 3*a and Video 5), we divide the landing process into three phases: running, lingering, and attaching. In the running phase (0-3.77 s), cells will swim and can perform roaming or orbiting. We note that misalignments between the flagellum and cell body axis tend to change the motion direction of the cell (*Figure 3*a, b). In the lingering phase (3.77-4.68 s), the cells demonstrate one of two states: a paused state or a tethered state, where the cell can move under the constraint of tethering pilus (see *Figure 3*a for the tethered state). At 3.77 s, one pilus attaches to the surface and acts as an anchor point to prevent the cell from moving away. Finally, in the attaching phase (≥ 4.68 s), cells remain on the surface motionless during the observation period most likely since they have effected irreversible attachment. Upon irreversible cell attachment, some of the free MSHA pili become attached to the surface firmly while others demonstrate fluctuations punctuated with intermittent attachment to the surface (Video 6).

**Figure 3.**
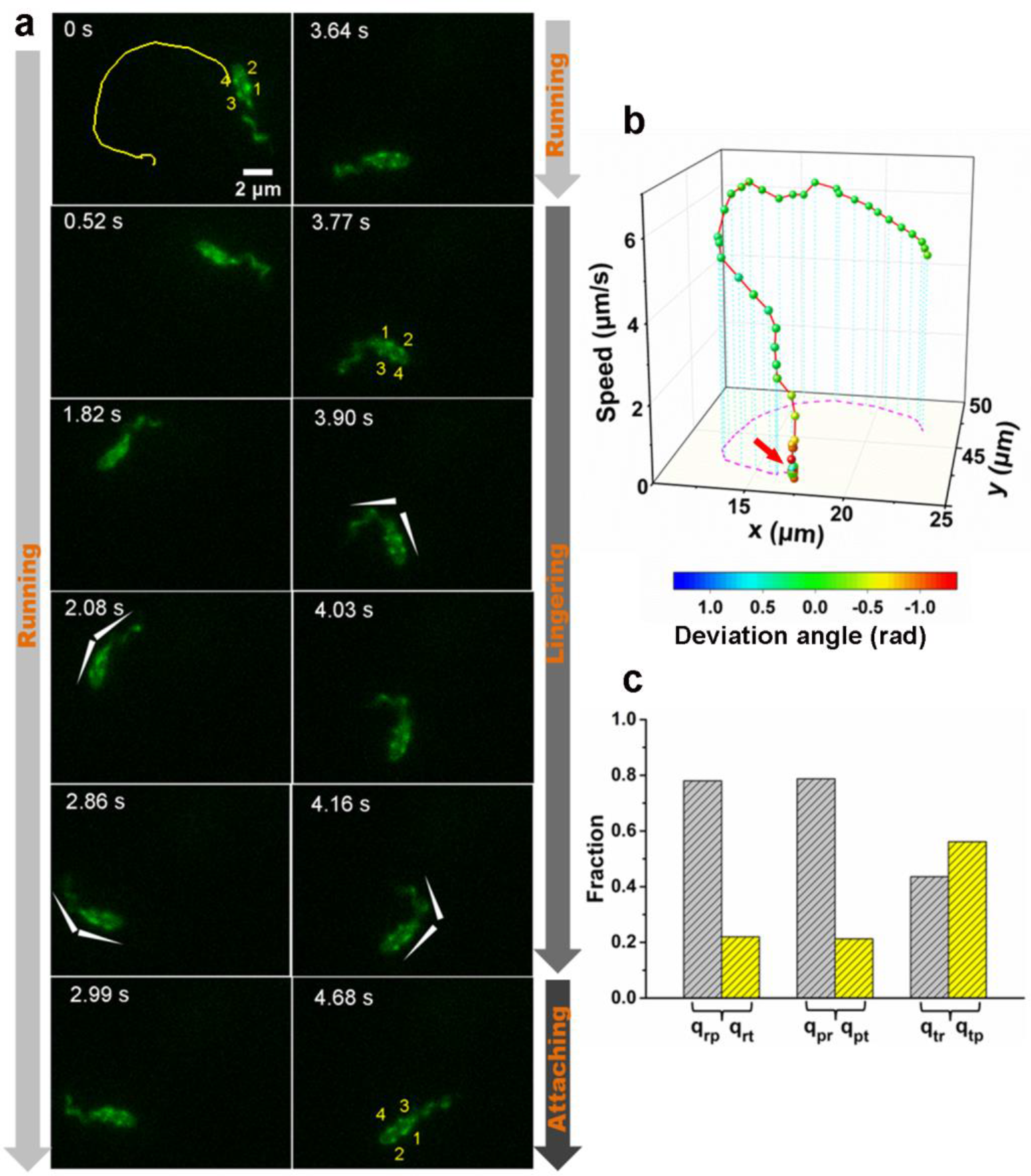
An example of a typical landing sequence of a *V. cholerae* cell with MSHA pili and flagella both labelled (MshAT70CFlaAA106CS107C). (a) Representative image sequences showing the behavior of MSHA pili and flagella. For easy identification, four pili of the example cell in Figure 3a were numbered from 1 to 4, which revolve around the major axis of the cell periodically as the cell swims. The white arrowheads indicated the orientation of cell body and flagellum. (b) A 3D plot of speed and deviation angle of the representative cell in panel (a) over its trajectory. The red arrow in panel (b) represents the position, where the pili touch surface, causing a deflection. (c) The conditional probabilities *q*_*ij*_ that the bacterium transitions from state *i* to *j*. The number of transition events used for estimating these conditional probabilities is 666. r: running state, t: tethered state, p: paused state. **Figure 3-source data 1. Figure 3b source data**. **Figure 3-source data 2. Figure 3c source data**.

During cell landing, transitions between the running and lingering phase, as well as between the two states of lingering phase are observed. The measured conditional probabilities *q*_*ij*_ that a cell transitions from state *i* to *j* show that the running phase has a relatively lower *q*_*rt*_ to the tethered state (∼22%) but a higher *q*_*rp*_ to the paused state (∼78%). Similarly, the paused state has a higher *q*_*pr*_ than *q*_*pt*_. In contrast, the tethered state shows similar *q*_*tr*_ and *q*_*tp*_, which are 45% and 55%, respectively (*Figure 3*c).

The single-cell dynamics in each specific phase/state is also characterized quantitatively. In the running phase of *V. cholerae*, we found that the period for body rotation is generally distributed between 0.25-2 s and is centered at ∼0.7 s (the rotation rate was ∼1.5 Hz) in LB+MC (*Figure 4*a). We measured the swimming speed, *v*, and the cell-body rotation rate, *ω*_c_, for each cell, and plotted *v* as a function of *ω*_c_ (see *Figure 4*b). By fitting the data, we found that *v* linearly increases with *ω*_c_ with a slope of |*v*/ω_c_| = 2.48±0.04 μm/radian.

**Figure 4.**
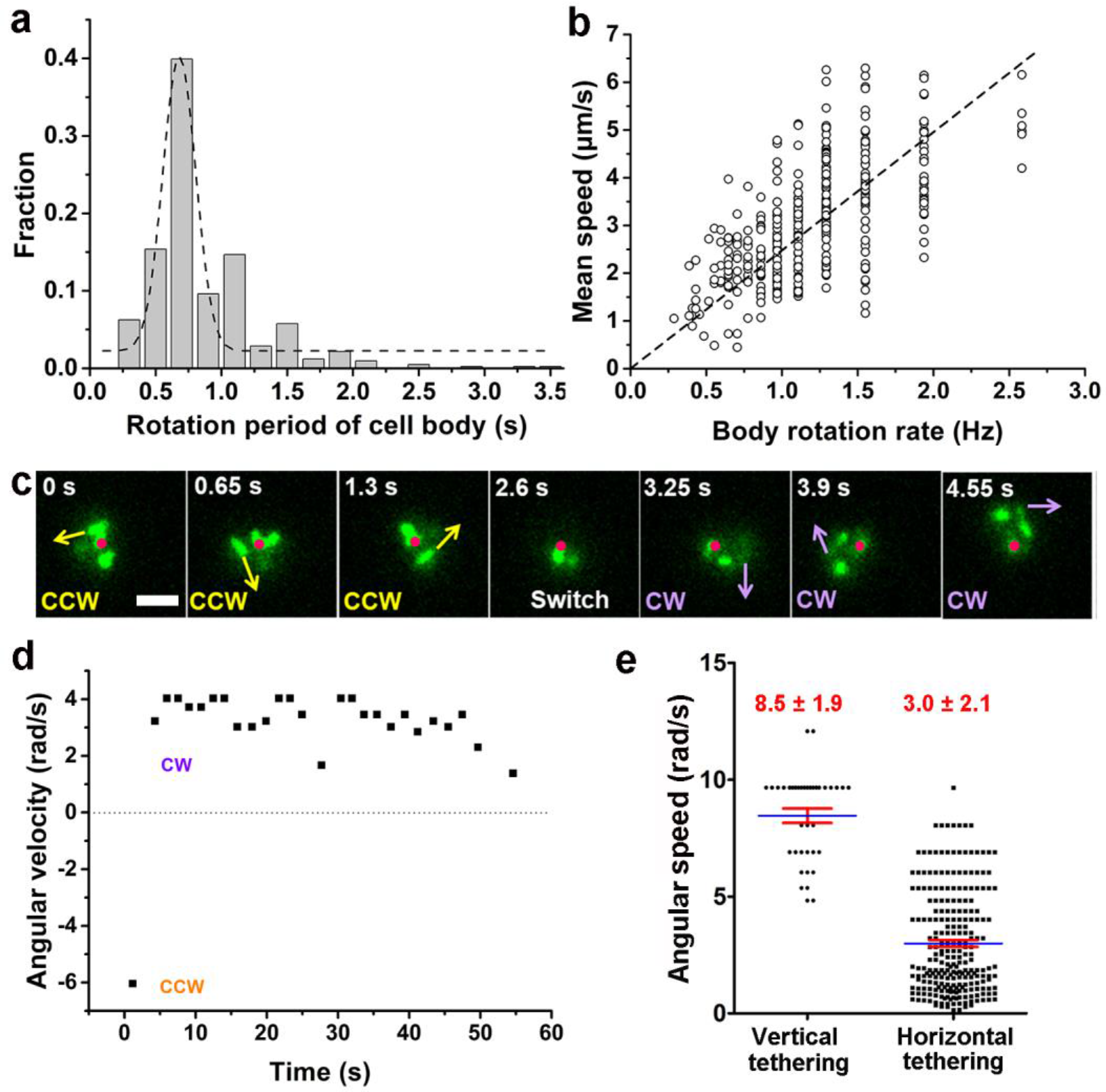
Characterization of running and tethered cells. (a) Distribution of the rotation period of cell body. The dashed line represents Gaussian fitting. A total of 416 rotation events from 54 cells were used for statistical analysis. (b) Measured relation between the rotation rate of cell body and the mean swimming speed of cell. The dotted line represents linear fitting result. N_cell_ = 47. (c) An example of a typical tethered motion, showing cell performing a circular motion around a center point (the red dot) with the direction of motion (noted by arrows) switched from CCW to CW. Scale bar, 2 μm. (d) The angular velocity of the tethered cell in panel (c) over a short duration showing a pair of CW (positive angular velocity) and CCW(negative angular velocity) intervals; (e) Distribution of angular speed of circular motion for horizontal (241 intervals from 25 cells) and vertical (38 intervals from 5 cells) tethered cells. **Figure 4-source data 1. Figure 4a source data**. **Figure 4-source data 2. Figure 4b source data**. **Figure 4-source data 3. Figure 4d source data**. **Figure 4-source data 4. Figure 4e source data**. **Figure 4-Figure supplement 1. Examples show positions of two poles and centroid of tethered motility**. The contributing MSHA pili were indicated by yellow arrows. (a) ∼1/2 position; (b) 1/3 or 2/3 position.

By contrast, a cell in the tethered state typically performs a circular motion around the attachment point (red dots in *Figure 4*c). The direction of the circular motion is also dynamic and can switch from counter-clockwise (CCW) to clockwise (CW) presumably due to a switch in the rotation direction of the flagellar motor (see 2.6 s, *Figure 4*c). Angular velocity is roughly constant during each circular-motion interval (i.e., in each CCW or CW period) and quickly changes sign after CCW-CW switching (*Figure 4*d and Video 7). Due to the distribution of pili across the cell body, tethering can occur at a pole or under the body, which leads to cells standing vertically or lying down horizontally to the surface, respectively. We find that standing tethered cells perform a faster circular motion (mean angular speed = 8.5 ± 1.9 rad/s) than lying ones (mean angular speed = 3.0 ± 2.1 rad/s) (*Figure 4*e). For the horizontal cells, different MSHA pili may be used to further anchor the cell to the surface. For example, two horizontally-tethered cells demonstrate different tethered-motion trajectories depending on the location of the anchoring MSHA pilus (*Figure 4-Figure supplement 1*). In addition to the fact that unattached pili may increase the likelihood that the cell will make irreversible attachment, we observe that MSHA pili appear to be able to attach to the surface along their entire length, and not just the tip (Video 8).

Interestingly, we find that the flagellum of attached cells frequently continues to rotate (Video 5, after 4.68 s), indicating that even after cell attachment, the flagellar motor is still active for some period. The flagellum will eventually stop rotating after a cell stay long enough on the surface (Video 9).

### Role of MSHA pili in cell landing is more apparent in viscoelastic non-Newtonian fluids than viscous Newtonian ones

To further investigate the dependence of MSHA pili function and hence cell landing on viscoelasticity, we compared cell motion behavior obtained in 2% LB, which is a Newtonian-fluid and has a viscosity ∼1 cP at 30 °C (*Utada et al., 2014*) and in 2% LB+1% MC, which is a non-Newtonian fluid and has a shear-dependent viscosity (*Figure 6*), for both WT and Δ*mshA* cells (*Figure 5* and *Figure 5-Figure supplement 1*).

**Figure 5.**
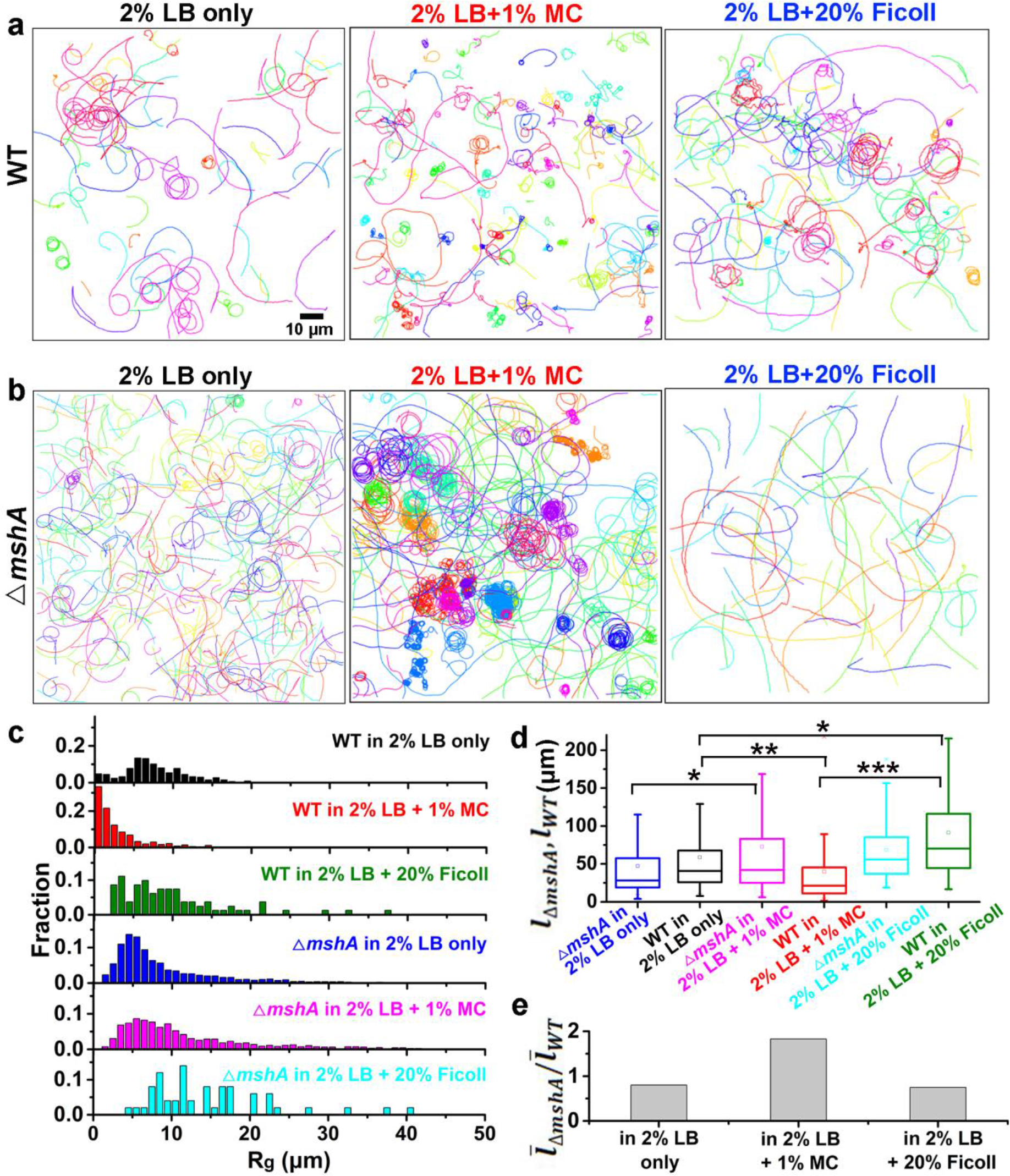
Role of MSHA pili in cell landing is more apparent in viscoelastic non-Newtonian solutions than viscous Newtonian ones. (a) Examples of WT cell trajectories showing both roaming and orbiting motilities in 2% LB only, 2% LB with 1% MC and 2% LB with 20% Ficoll; (b) Examples of cell trajectories of Δ*mshA*; (c) Histograms of R_g_ of WT and Δ*mshA* in different solutions; (d) A box plot summary of path lengths of WT and Δ*mshA*; (e) The ratio of mean path length between Δ*mshA* and WT,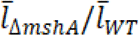. **Figure 5-source data 1. Figure 5c source data**. **Figure 5-source data 2. Figure 5d source data**. **Figure 5-source data 3. Figure 5e source data**. **Figure 5-Figure supplement 1. Motility characterization of WT and Δ*mshA* cells in 2% LB only and in 2% LB with 1% MC**. (a) Histograms of deviation angle for WT in 2% LB only. (b) Histograms of deviation angle for WT in 2% LB+1% MC viscous solution. Black represents orbiting motility and red represents roaming motility. (c-d) Mean square displacements (MSDs) of WT (c), Δ*mshA* (d) in 2% LB only and 2% LB+1% MC viscous solution.

**Figure 6.**
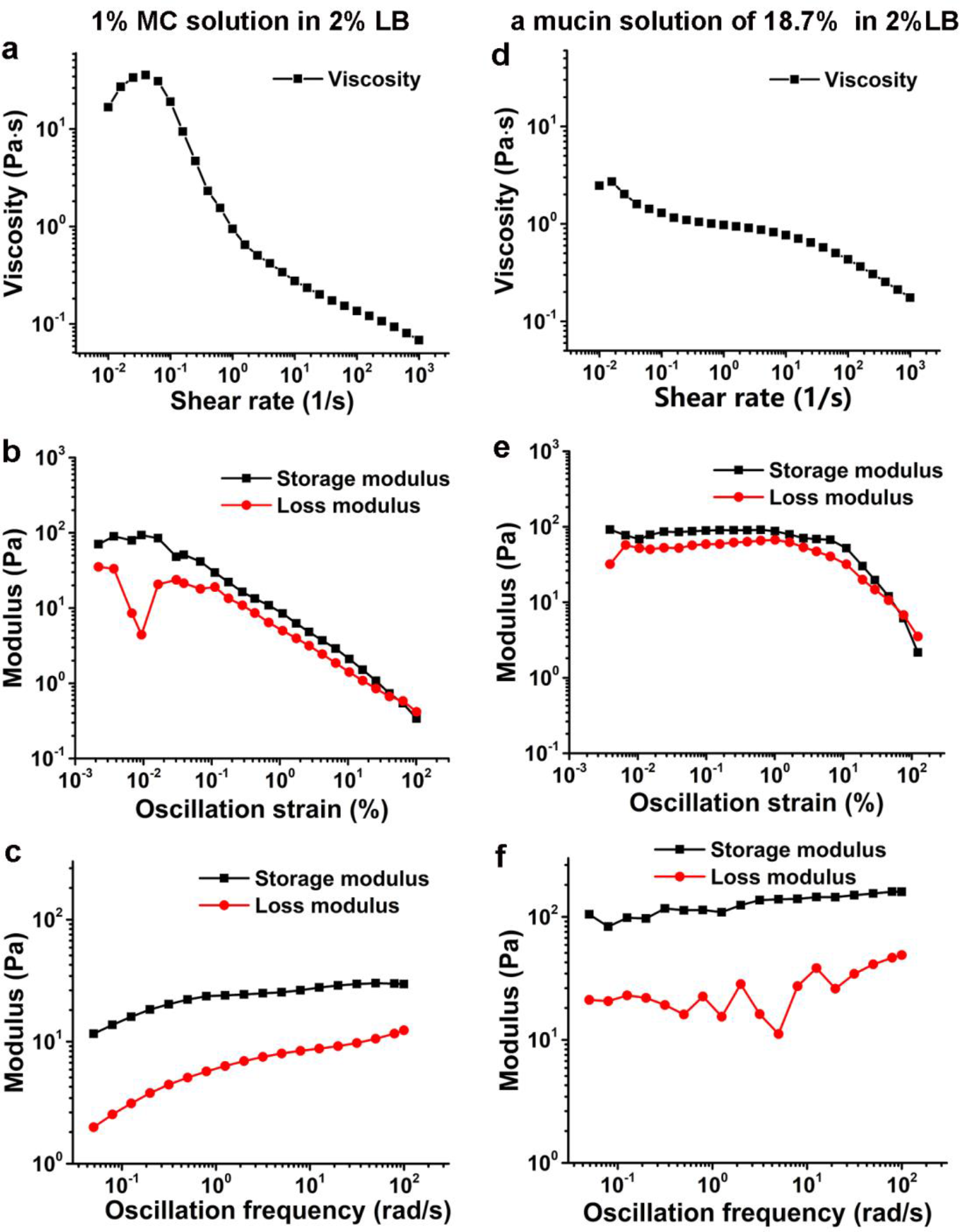
Viscoelasticity characterization of 1% MC solution in 2% LB (a-c) and a mucin solution of 18.7%(w/w) in 2% LB (d-f) at 26°C. (a) and (d) show viscosity as a function of shear rate. (b) and (e) display modulus as a function of the oscillation strain, using cone-plate geometry. (c) and (f) show modulus as a function of the oscillation frequency under an oscillation strain of 0.1%, using cone-plate geometry. **Figure 6-source data 1. Figure 6a source data**. **Figure 6-source data 2. Figure 6b source data**. **Figure 6-source data 3. Figure 6c source data**. **Figure 6-source data 4. Figure 6d source data**. **Figure 6-source data 5. Figure 6e source data**. **Figure 6-source data 6. Figure 6f source data**.

We observed WT cells to demonstrate roaming and orbiting motilities in both solutions (*Figure 5*a). The histograms of deviation angle of each type of motility obtained in the two solutions are also similar (*Figure 5-Figure supplement* 1a and b). These results indicate that roaming and orbiting motilities of cells are robust against the tested viscoelastic environment. Although the general motility pattern is similar in both solutions, the motion of cells, as expected, is slowed significantly in LB+MC. The average speed of WT cells for near-surface motion is reduced by ∼ 22 times from 86.7 ± 32.9 μm/s (mean ± standard deviation) in 2% LB to 3.8 ± 2.6 μm/s in LB+MC. Similarly, the average speed of Δ*mshA* cells is also decreased by ∼ 12 times from 80.0 ± 15.0 μm/s in 2% LB to 6.5 ± 1.4 μm/s in LB+MC. The slow-down of motion can also be seen clearly from their mean square displacement curves (*Figure 5-Figure supplement* 1c, d), which show similar shape but very different time scales.

However, WT and Δ*mshA* cells also show differences in their motility behavior in these two solutions(*Figure 5*). In LB+MC, WT cells tend to land on the surface soon after approaching it (less than one round in orbiting motility) and more tethered motions are observed, which leads to more irregular and tortuous trajectories and smaller R_g_ for WT cells compared with the case of 2% LB (*Figure 5*c). By contrast, Δ*mshA* cells show very similar R_g_ distributions in the two types of solutions (*Figure 5*c). More interestingly, compared with WT, in LB+MC, a large proportion of Δ*mshA* cells show orbiting for a substantial large number of cycles, as shown in Figure 5b. Quantitatively, this can be seen by the calculated mean path length, 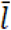, which is 39.7 ± 51.2 μm for WT and 72.5 ± 99.1 μm for Δ*mshA* in LB+MC, whereas the corresponding value in 2% LB is 58.7 ± 63.1 μm for WT and 47.2 ± 50.8 μm for Δ*mshA*. To see how the role of MSHA pili varies with viscoelasticity, we can calculate the ratio of mean path length between Δ*mshA* and WT,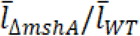, for each type of solution, which is ∼1.8 in LB+MC and ∼ 0.8 in 2% LB only, respectively (*Figure 5*d and e). So loss of MSHA pili results in a more dramatic increase in mean path length in LB+MC than in 2% LB. Moreover, such prolonged orbiting motions of Δ*mshA* were not observed in a 20% Ficoll solution (*Figure 5*), which shows a high viscosity but still belongs to a Newtonian fluid (*Winet, 1976*). As shown in Figure 5, the cell motility behaviors in 20% Ficoll are similar to those in 2% LB only except that the average cell speed for near-surface motion is dramatically slowed in 20% Ficoll, which is 9.3 ± 4.3 μm/s for WT and 10.0 ± 3.5 μm/s for Δ*mshA*. The cell trajectories in 20% Ficoll are similar to those in 2% LB only and consequently, the ratio of 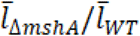 is ∼0.75 in 20% Ficoll, very close to ∼0.8 in 2% LB only. Together, these results indicate that the elastic properties of viscoelastic solutions can also affect cell motility behavior, and the role of MSHA as a braking and anchoring machine in cell landing is more apparent in viscoelastic non-Newtonian fluids than viscous Newtonian fluids.

## Discussion

The first step in *V. cholerae* biofilm formation is the transition from planktonic swimmers to stationary surface attached cells; this process is mediated by the landing process (*Teschler et al., 2015*). In this study, the combination of cell appendage labelling with high-resolution spatio-temporal imaging allows us to quantitatively deconstruct the landing process into three stages: running, lingering, and attaching, as summarized in *Figure 7*. During the running phase, cell motion is powered by flagellar rotation, which simultaneously induces a counter-rotation of cell body. When swimming cells come to within a distance that is comparable to the length of a typical pilus from a surface, dangling pili may brush against the surface, thereby deflecting the trajectory. Typical MSHA pili are ∼0.4-1.2 μm in length. During near surface swimming, cell body rotation actively brings MSHA pili into close proximity with the underlying surface where friction between pili and the surface can slow the cells, or, transient adhesions can be made, which may even arrest cell motion. Here, we make an analogy to the slow-down and stop effected by the brake system of a car. During near-surface swimming, it has been suggested that hydrodynamic forces cause the cell bodies of swimming rod-like bacteria take on a tilted, non-parallel, orientation to the surface (*Vigeant et al., 2002*). In the case of *P. aeruginosa*, whose TFP are distributed with a strong bias toward a particular pole (*Skerker et al., 2001*), pili-surface contact will depend on which pole is closer to the surface. In contrast, the homogeneous distribution of MSHA pili on *V. cholerae* (see *Figure 1c*) may be more efficient at slowing such tilted cell bodies by increasing the probability that pili encounter the surface.

**Figure 7.**
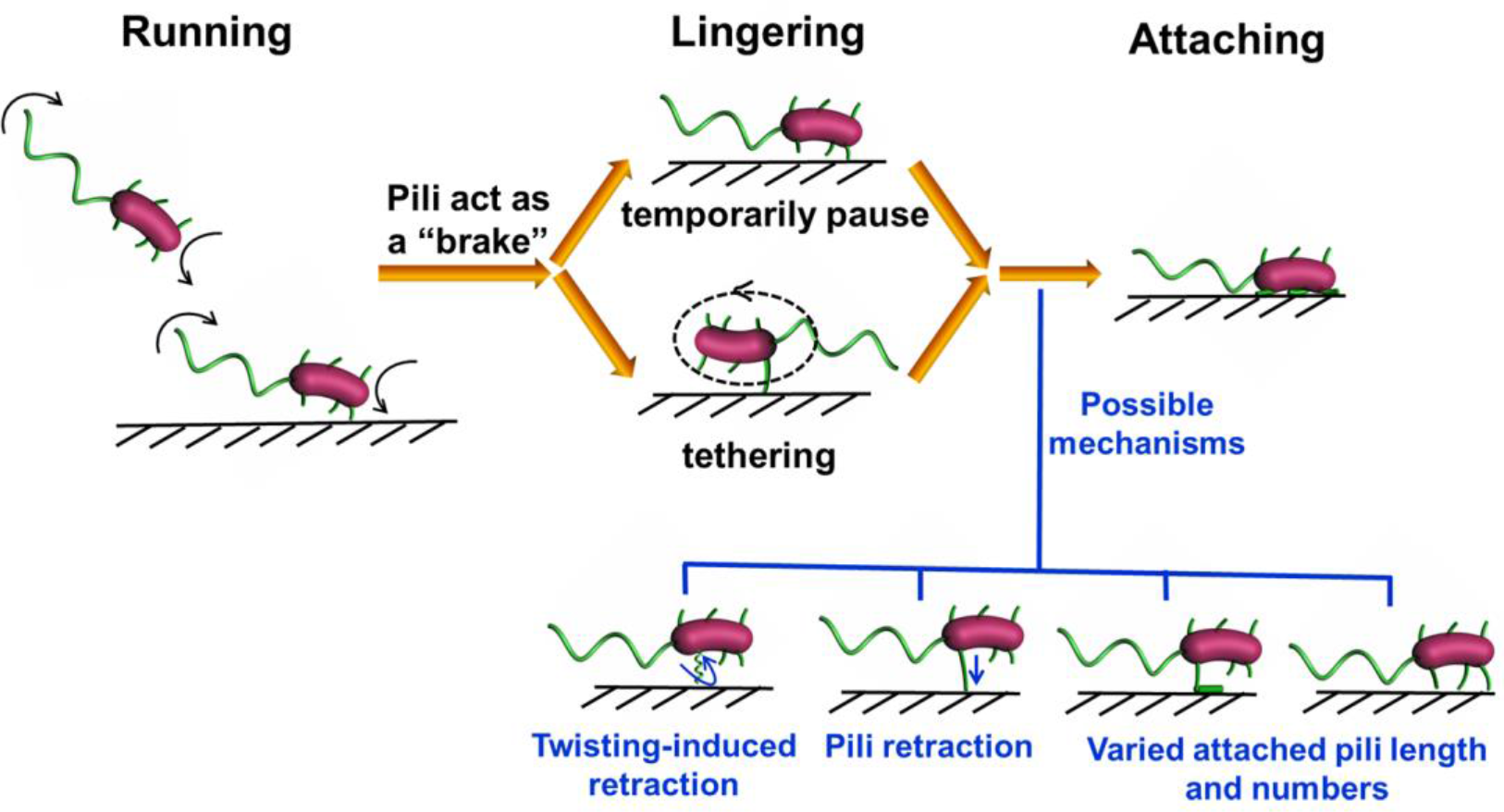
Diagrammatic sketch for the landing process of *V. cholera* cells on a substrate.

If the contact-induced adhesion between MSHA pili and the surface is sufficiently strong to arrest forward motion, the cell will either pause or commence tethered motion centered about the point-of-adhesion. Cells rotating at an angle closer to the surface have a slower angular velocity (*Figure 4e*), to which hydrodynamic effect presumably has an important contribution (Bennett et al., 2016). This suggests that for cells demonstrating tethered motion, a progressive twisting of the surface-attached pilus fiber during the circular motion of cells may gradually cause the circular motion to stop by pulling the cell body ever closer to the surface. Although twitching has not be observed in *V. cholerae*, this is one mechanism by which retraction-like dynamics may be achieved (*Charles et al., 2019*), possibly in tandem with actual retraction of MSHA pili, which has been shown recently in a different strain of *V. cholera* (*Floyd et al., 2020*). Under our conditions, we have not observed MSHA pili retraction events nor have we seen bacterial cells that gradually acquire fluorescence when only maleimide dyes were used. These results are consistent since in bacteria where pilus retraction does occur, such as in the TAD pili of *Caulobacter crescentus* (*Ellison et al., 2017a*), ChiRP pili of *V. cholera* (*Ellison et al., 2018*), and TFP of *P. aeruginosa* (*Skerker et al., 2001*), the cell body gradually becomes fluorescent due to internalization of labeled pili by retraction. Such phenotypical differences may be due to the different experimental conditions used in each study and require more work to fully elucidate.

In addition to possible hydrodynamical effects, our observation that MSHA pili are able to adhere to surfaces along their entire length highlights their versatility and likely increases the chances of the formation of a cell-surface attachment. The ability to adhere not only at the distal tip, contrasts with the TFP of *P. aeruginosa* (*Skerker et al., 2001*) and ChiRP pili of *V. cholerae* (*Ellison et al., 2018*) who show the pilus-subject interactions mainly mediated by the pilus tip. Thus, for *V. cholerae*, the strength of adhesion between a cell and a surface that is mediated by an individual MSHA pilus appears to be more complicated to model with a single point of attachment. Rather, cells can enhance the adhesion strength by increasing both the length and the number of the MSHA pilus adhered to the surface. This will facilitate cells to become irreversibly attached.

Similar running and lingering phases for cells near surface motion has also been reported in enterohaemorrhagic *E. coli* (EHEC) cells (*Perez Ipiña et al., 2019*), where results suggested that by choosing the optimal transition rates, EHEC bacterial diffusivity is maximized and the surface exploration efficiency is greatly improved. In a future work, it will be interesting to apply similar analysis in *V. cholerae*.

In this study, the data collection of *V. cholerae* cells was performed mainly in the solution of 2% LB+1% MC. Rheological measurements show that its viscosity is shear-dependent and has a higher storage modulus than loss modulus in the tested oscillation frequency range, thus it is a non-Newtonian fluid (*Figure 6a-c*). On the other hand, the rheological measurements of a mucin solution in 2% LB with a mucin concentration of 18.7% (w/w), which was obtained by measuring the concentration of mixtures scraped from the fresh mouse intestine surfaces, show similar viscoelastic behavior with a higher storage modulus than loss modulus in the same tested oscillation frequency range (*Figure 6d-f*). So compared with Newtonian Ficoll solutions, LB+MC is better to simulate the viscoelastic environment that *V. cholerae* cells encounter in the mucus layer of animal intestines. In such viscoelastic environments, Millet *et al*. (*Millet et al., 2014*) observed considerable differences of bacterial localization in different parts of small intestine and found that *V. cholerae* motility exhibits a regiospecific influence on colonization, indicating viscoelastic intestinal mucin is a key factor limiting colonization. To directly observe cell motion in mucus solutions made from lyophilized mucus powders or fresh mucus-containing solutions scraped from the surfaces of mouse intestines is technically challenging due to the complicated inhomogeneous environment with too many impurities (data not shown). In this work, by direct visualization of pili and flagellum of cells during their landing process in LB+MC that mimic the real mucus solutions, we find that *V. cholerae* cells can move well in this viscoelastic solution under our conditions. Moreover, we show that the effect of MSHA pili as a braking and anchoring machine on cell landing is more apparent in LB+MC than in 2% LB only or 20% Ficoll solutions, suggesting that MSHA pili might play an even more important role for cell surface attachment in viscoelastic non-Newtonian environments such as in the mucus layer of small intestines.

To summarize, in this work, using fluorescence imaging with labeled pili and flagellum, we show a comprehensive picture of the landing dynamics of *V. cholerae* cells in viscoelastic environments and provide a direct observational evidence exhibiting the role of MSHA pili during cell landing. We hope this can shed insights into the prevention and control of *V. cholerae* infections.

## Materials and methods

### Bacterial strains

Bacterial strains, plasmids, and primers used in this study are listed in Table 1. *V. cholerae* El Tor C6706 (*Joelsson et al., 2006*) was used as a parental strain in this study. C6706 and mutants were grown at 30 °C or 37 °C in Luria-Bertani (LB) supplemented with 100 µg/mL streptomycin, 50 µg/mL kanamycin, 1 µg/mL chloromycetin where appropriate. *E. coli* strains harboring plasmids were grown at 37 °C in LB supplemented with 100 µg/mL ampicillin. The optical densities of bacterial cultures were measured at 600 nm (OD_600_) using a UV-vis spectrophotometer.

**Table 1.** Strains, plasmids, and primers used in this study

| Strain | Description | Source or reference |
| --- | --- | --- |
| parent | C6706 Sm<sup>R</sup> | (Joelsson *et al.*, 2006) |
| $\Delta mshA$ | C6706 Sm<sup>R</sup>, VC1807::Cm<sup>R</sup>, *mshA* knockout | This study |
| $\Delta flaA$ | C6706 Sm<sup>R</sup>, VC1807::Cm<sup>R</sup>, *flaA* knockout | This study |
| MshA<sup>T70C</sup> | C6706 Sm<sup>R</sup>, VC1807::Km<sup>R</sup>, MshAT70C | (Ellison *et al.*, 2017b) |
| FlaA<sup>A106C</sup> | C6706 Sm<sup>R</sup>, VC1807::Cm<sup>R</sup>, FlaAA106C | This study |
| FlaA<sup>S107C</sup> | C6706 Sm<sup>R</sup>, VC1807::Cm<sup>R</sup>, FlaAS107C | This study |
| FlaA<sup>A106CS107C</sup> | C6706 Sm<sup>R</sup>, VC1807::Cm<sup>R</sup>, FlaAA106CS107C | This study |
| FlaA<sup>E332C</sup> | C6706 Sm<sup>R</sup>, VC1807::Cm<sup>R</sup>, FlaAE332C | This study |
| FlaA<sup>G23C</sup> | C6706 Sm<sup>R</sup>, VC1807::Cm<sup>R</sup>, FlaAG23C | This study |
| FlaA<sup>N26C</sup> | C6706 Sm<sup>R</sup>, VC1807::Cm<sup>R</sup>, FlaAN26C | This study |
| FlaA<sup>N83C</sup> | C6706 Sm<sup>R</sup>, VC1807::Cm<sup>R</sup>, FlaAN83C | This study |
| FlaA<sup>S325C</sup> | C6706 Sm<sup>R</sup>, VC1807::Cm<sup>R</sup>, FlaAS325C | This study |
| FlaA<sup>S87C</sup> | C6706 Sm<sup>R</sup>, VC1807::Cm<sup>R</sup>, FlaAS87C | This study |
| FlaA<sup>S376C</sup> | C6706 Sm<sup>R</sup>, VC1807::Cm<sup>R</sup>, FlaAS376C | This study |
| FlaA<sup>V117C</sup> | C6706 Sm<sup>R</sup>, VC1807::Cm<sup>R</sup>, FlaAV117C | This study |
| MshA<sup>T70C</sup>, $\Delta flaA$ | C6706 Sm<sup>R</sup>, VC1807::Km<sup>R</sup>, *flaA* knockout | This study |
| MshA<sup>T70C</sup>, FlaA<sup>A106C</sup> | C6706 Sm<sup>R</sup>, VC1807::Km<sup>R</sup>, FlaAA106C | This study |
| MshA<sup>T70C</sup>, FlaA<sup>S107C</sup> | C6706 Sm<sup>R</sup>, VC1807::Km<sup>R</sup>, FlaAS107C | This study |
| MshA<sup>T70C</sup>, FlaA<sup>A106CS107</sup> | C6706 Sm<sup>R</sup>, VC1807::Km<sup>R</sup>, FlaAA106CS107C | This study |

### Flagellin and pilin mutagenesis

Following the protocol in Ellison *et al*. (*Ellison et al., 2019; Ellison et al., 2017a*), we first predicted 10 amino acid residues in *V. cholerae* flagellin FlaA for cysteine replacement. Then the *flaA* knockout and FlaA sequences containing the FlaAA106C, FlaAS107C, FlaAA106CS107C, FlaAE332C, FlaAG23C, FlaAN26C, FlaAN83C, FlaAS325C, FlaAS87C, FlaAS376C, FlaAV117C knock-in were constructed using the MuGENT method (*Dalia et al., 2014*). The FlaAA106C, FlaAS107C, and FlaAA106CS107C knock-in were constructed by cloning the fragment into the suicide vector pWM91 containing a *sacB* counter-selectable marker (*Metcalf et al., 1996*). The plasmids were introduced into *V. cholerae* by conjugation and mutations were selected for double homologous recombination events. The MshAT70C mutation can be successfully labeled with thiol-reactive maleimide dyes has been described previously (*Ellison et al., 2017a*), and MshAT70C was constructed using the MuGENT method to light MSHA pilus. All mutants were confirmed by DNA sequencing.

### Hemagglutination assays

Mannose-sensitive hemagglutination by *V. cholerae* was measured as described previously (*Gardel et al., 1996*). Briefly, bacteria were grown to the mid-logarithmic phase in LB medium. Initial concentrations of approximately 10^10^ CFU/mL were two-fold diluted with KRT buffer in U-bottomed wells of 96-sample microtiter dishes. Sheep erythrocytes were washed in PBS and resuspended in KRT buffer for a final concentration of 10% vol/vol. Equivoluminal erythrocyte were added into serially diluted bacterial suspensions and the plates were gently agitated at room temperature for 1 min. Samples were checked for hemagglutination after 2 h at room temperature (RT).

The results of the Hemagglutination assay test show that MshAT70C displays similar behavior to WT, which indicates that the point mutation in MSHA does not affect MSHA pilus function (*Figure 1-Figure supplement 4*).

### Cell tracking and analysis

#### Preparation of viscous solutions

To change the solution viscosity, methyl cellulose (MC) (M20, 4000 cp, Solarbio, China) solutions were prepared by dissolving 0% and 1% (wt/vol) MC in 2% Luria-Bertani (LB) motility medium (containing 171 mM NaCl). 20%(w/v) Ficoll 400 (MW=400 kDa, Yuanye Bio-Technology Co.,Ltd, China) dissolved in 2% LB medium was also prepared for a control experiment.

#### Preparation of the small intestinal mucus samples from ICR female mice

Six-week-old female ICR mice were provided with drinking water with 10 g/L streptomycin and 0.2g/L aspartame for two days. To clean the small intestine content, food will be removed before 24 h. Mice were sacrificed, the small intestine was cut open with sterile scissors and the mucus layer was gently extracted with a cell scraper. Samples were weighed to get wet weight first, then frozen overnight at -80 °C. After that, they were freeze-dried using a freeze-dryer (Labconco, the United States) in -40 °C low temperature vacuum. Next, these dried samples were weighted again to get dry weight. The obtained average value of dry/wet ratio from three repeats is 18.7%.

#### Cell imaging

For the *V. cholerae* motility observation in 2% LB without MC, overnight cultures in LB were resuspended and diluted with 2% LB to an OD_600_ ranging from 0.01-0.03. Then the bacterial suspension was injected into a flow cell, which contained the same media. Imaging was performed using a Phantom V2512 high-speed camera (Vision Research, USA) collecting ∼200 000 bright-field images at 5 ms resolution with a 100× oil objective on a Leica DMi8 inverted microscope (Leica, Germany) at a set temperature value of 30°C.

For the *V. cholerae* motility observation in 2% LB with 1% MC (henceforth, this medium is referred to LB+MC), overnight cultures in LB were resuspended and diluted with LB+MC to a final OD_600_ of 0.01-0.03. Then, the bacteria were incubated at 37°C for 20 min to allow them to adapt to the new environment and were then used immediately. Bacteria samples were pipetted onto standard microscope slides with a 8 mm diameter spot and then were sealed with a coverslip using a 1 mm thick secure spacer. Imaging was performed using EMCCD camera (Andor iXon Ultra 888) collecting ∼10 000 bright-field images with a time resolution of 90ms at a set temperature value of 30°C. Similar protocols were carried out for observations in 20% Ficoll solutions and in the small intestinal mucus samples of mice.

#### Cell-tracking algorithms and analysis

The images were preprocessed using a combination of software and algorithms adapted from the methods described (*Lee et al., 2016; Utada et al., 2014; Zhao et al., 2013*) and written in MATLAB R2015a (Mathworks) by subtracting the background, scaling, smoothing and thresholding. After image processing in this way, the bacteria appear as bright regions. The bacteria shape was fit with a spherocylinder. Then the geometric information of the cell, such as location of the centroid and two poles, and the length and width of the bacterium were collected. Trajectory reconstruction was also achieved for further analysis.

The motility parameters (*Utada et al., 2014*), such as instantaneous speed, deviation angle, radius of gyration (R_g_), MSD and mean path length 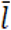 were calculated to further characterize the near-surface motility of *V. cholerae*. The instantaneous speed was calculated via |*r*_i+1_-*r*_i_|/Δ*t*, where *r*_i_ is the cell position vector in frame *i* and Δ*t* is the time interval between two consecutive frames. The deviation angle of cell motion is defined as the angle between its cell body axis and the direction of motion. The radius of gyration, *R*_*g*_, is a statistical measure of the spatial extent of the domain of motion given by an ensemble of points that define a trajectory(*Rubenstein, 2003*). The square of this quantity is defined as 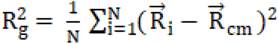, where *N* is the number of points in the tracked trajectory, 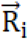 is the position vector corresponding to the *i*-th point on the trajectory, 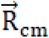 is the position vector of the center-of-mass. The MSD of cells was calculated via ⟨Δ*r*^2^(τ) ⟩= ⟨[*r*(*t* + *τ*) − *r*(*t*)] ^2^⟩, where *r*(*t*) is the position vector of a cell at time *t*, and *τ* represents the time lag. The MSD provides information on the average displacement between points in the motility trajectory separated by a fixed time lag. Mean path length 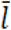 were calculated as the average of the total travelling distance of each tracked cell in the field of view.

#### MSHA pilus labeling, imaging, and quantification

Pilin labeling was achieved using Alexa Fluor 488 C5 Maleimide (AF488-mal; ThermoFisher Scientific, cat. no. A10254) or Alexa Fluor 546 C5 Maleimide (AF546-mal; ThermoFisher Scientific, cat. no. A10258), which were dissolved in DMSO, aliquoted, and stored at -20°C while being protected from light.

*V. cholerae* cultures were grown to mid-log phase (OD_600_ 0.8-1.5) before labeling. ∼100 µL of culture was mixed with dye at a final concentration of 25 µg/mL (*Ellison et al., 2017a*) and incubated at RT for 5 min in the dark. Labeled cultures were harvested by centrifugation (900×g, 5 min) and washed twice with PBS, resuspended in 200 µL PBS and imaged immediately. Images were collected using an EMCCD camera on a Leica DMi8 inverted microscope equipped with an Adaptive Focus Control system. The fluorescence of cells labeled with AF488-mal and AF546-mal were detected with FITC and Rhod filter, respectively. The cell bodies were imaged using phase contrast microscopy.

To quantify the number of MSHA pili per cell and cell length, imaging was done under 0.2% PBS gellan gum pads. The cell lengths were measured using ImageJ.

We used AF546-mal and AF488-mal, in turn, for the two-color labeling to observe the growth of pili. We first, labeled log-phase cells with AF546-mal for the primary staining by incubating for 20 min, followed by two successive washes in PBS by centrifugation. The cells were then resuspended in LB and incubated for an additional 40 min at 30 °C. For the secondary staining, we incubated the cells in AF488-mal for 5 min, washed twice with PBS, and then imaged the cells immediately using phase contrast, FITC, and RhoD channels.

#### Fluorescence video acquisition of MSHA pilus-labelled cells motility in LB + MC

The labeled cells were centrifugated, resuspended in ∼20 µL PBS, and then diluted in 500 µL of the viscous solution of LB + MC. The solution was then immediately pipetted onto a standard microscope slides. Fluorescence images were acquired at 130 ms intervals for a total of about 2-5 min. After a few minutes of fluorescence imaging, most cells in the field of view have attached to the surface, while the fluorescence was bleached due to the continuous exposure. We recorded images from different locations to capture new instances of bacterial movement and adhesion events.

#### Rheological measurements

Rheological measurements of 1% MC+2% LB solution and 18.7% (w/w) mucin (Sigma, USA) in 2% LB solution were carried out on a rheometer (DHR-2, TA Instruments, Waters LLC) using a cone-plate geometry with a diameter of 40 mm and a cone angle of 5° at 26°C. Here, because when they were observed on a microscope, the real temperature of samples at the observation window site is lower than the set value of 30°C due to the relatively low thermal conductivity of glass, which is estimated to be 26 °C ∼28 °C, the rheological measurements were performed at 26 °C. The viscosity curves were determined at shear rates of 0.01-1000/s. And the storage modulus and loss modulus as a function of the testing oscillation strain and oscillation frequency were also recorded. For each tested solution, it was left standing for 15 min prior to each measurement for equilibrium, and it was covered with a thin layer of silicon oil to prevent loss of moisture during measurement.

### Statistical analysis

Statistical significance was determined with One way ANOVA followed by Tukey’s multiple comparison test comparing the different groups. (*p < 0.05; **p < 0.01; ***p < 0.001). The data were analyzed using the Prism 5.0 software program (GraphPad Software, La Jolla, CA, USA).

## Acknowledgements

We thank Zhanglin Hou and Thomas G. Mason for their help with scientific discussions.

## Additional information

### Funding

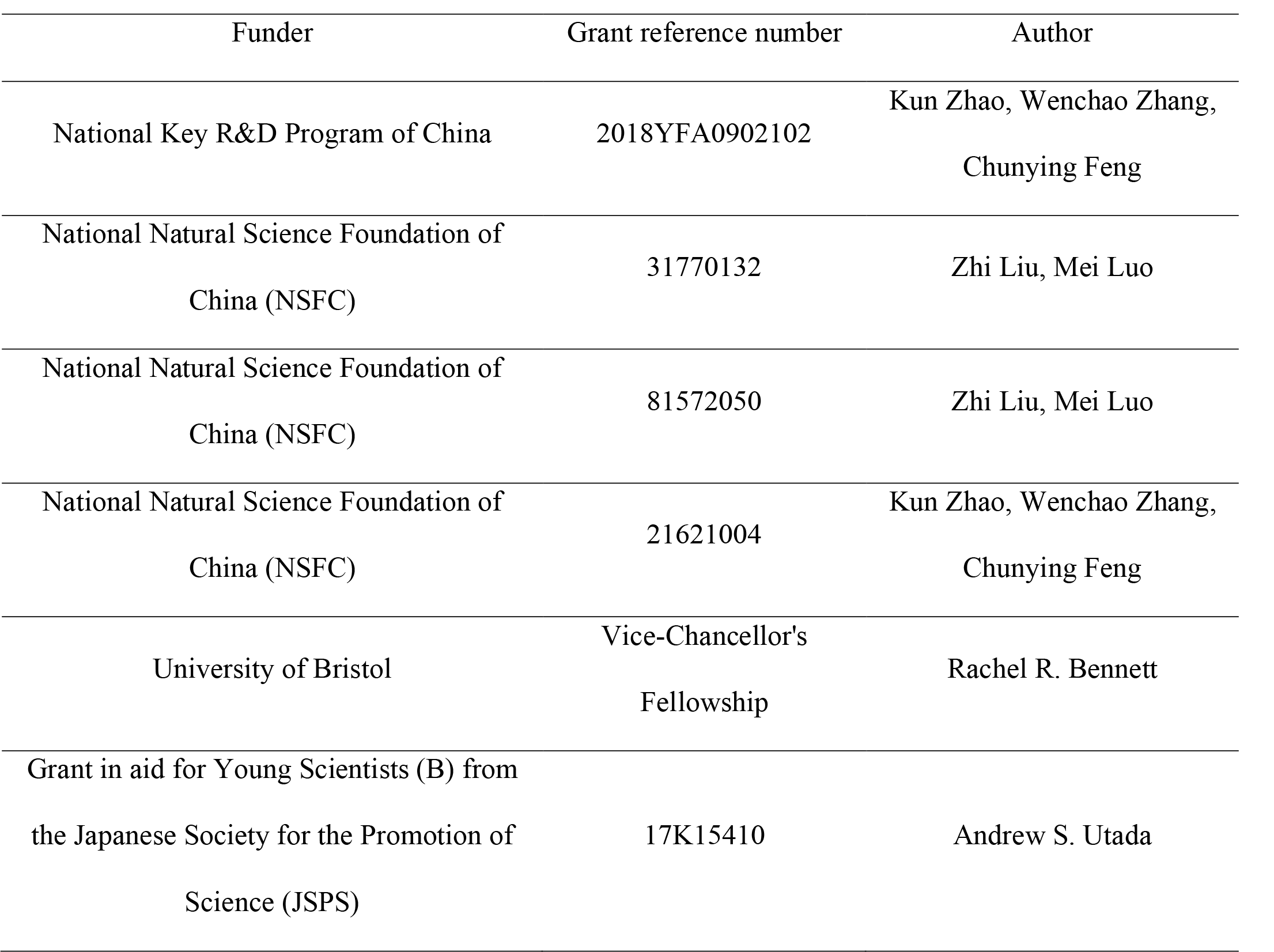

The funders had no role in study design, data collection and interpretation, or the decision to submit the work for publication.

## Additional files

Supplementary File 1. Including Supplementary table S1 and Supplementary references.

Source code 1. Source code for Figure 5a and b trajectory plots.

Transparent reporting form

## Data availability

Source data files and/or MATLAB code have been provided for Figures 1-6.

## Video Legends

**Video 1. Time-lapse fluorescence imaging showing a typical roaming cell (indicated by the arrowhead) with labeled MSHA pili in 2% LB+1% MC viscous medium**. This video was shown every 390 ms for 98 s and displayed at 20 frames per second (fps).

**Video 2. Time-lapse fluorescence imaging showing a typical orbiting cell with labeled MSHA pili in 2% LB+1% MC viscous medium**. This video was recorded every 130 ms for 15 s and displayed at 10 fps.

**Video 3. Time-lapse fluorescence imaging showing switch of pili**. When transient pauses happened, the attached pilus could be switched from one to another or more. See also Figure S4. This video was recorded every 70 ms for 10 s and displayed at 5 fps.

**Video 4. Time-lapse fluorescence imaging showing linear motion bent into circular motion that is centered around the attachment point between MSHA pili and the surface, which can act as an anchor point**. This video was recorded every 130 ms for 8 s and displayed at 10 fps.

**Video 5. Time-lapse fluorescence imaging showing dynamic movements of a MshAT70CFlaAA106CS107C cell with labeled flagellum and MSHA pili in 2% LB+1% MC viscous medium**. This video was recorded every 130 ms for 13 s and displayed at 10 fps.

**Video 6. Time-lapse fluorescence imaging showing five MSHA pili of a WT cell stuck to the surface and kept still or fluctuated frequently**. This video was recorded every 460 ms for 25 s and displayed at 10 fps.

**Video 7. Time-lapse fluorescence imaging showing a typical tethered cell performing a circular motion around a fixed point with the direction of motion switched from CCW to CW**. See also Figure 4c. This video was recorded every 130 ms for 6 s and displayed at 5 fps.

**Video 8. Time-lapse fluorescence imaging showing different adhesion points of a pilus**. When the tip of the pilus was free (∼3.5 s), the upper part of the pilus was still capable of keeping the cell adhered. This video was recorded every 130 ms for 13 s and displayed at 10 fps.

**Video 9. Time-lapse fluorescence imaging showing the motion evolution of flagellum from rotating to stopping eventually**. This video was recorded every 130 ms for 10 s and displayed at 10 fps.

