## Supplementary information for "Crash landing of *Vibrio cholerae* by MSHA pili-assisted braking and anchoring in a viscoelastic environment"

**This file includes:**

Tables S1

Supplementary References

### Table S1. Plasmids, and primers used in this study.

| Plasmids | Description | Source or reference |
|---|---|---|
| pWM91 | Suicide vector | (*Metcalf et al., 1996*) |
| pML3 | pWM-FlaAA106C | This study |
| pML4 | pWM-FlaAS107C | This study |
| pML5 | pWM-FlaAA106CS107C | This study |

| Primer Name | Primer Sequence (5'→ 3') | Description |
|---|---|---|
| VC0409-F1 | CTTGTATGGCGCACTCAACG | *mshA* knockout |
| VC0409-R1-3S | CAGCGCTAATTCAGTTTAAGCGGCCATAGCTACGCAGCATTACTGCAAGG | *mshA* knockout |
| VC0409-F2-3S | GCTATGGCCGCTTAAACTGAATTAGCGCTGCGTTATACAGCTGCAACCTC | *mshA* knockout |
| VC0409-R2 | CAAGCATAGCCTTGCTGTTC | *mshA* knockout |
| VC0409-Mut-T70C-R1 | GTCTAAACATTCAATGCCTTTAATTGCAGCTCGTCC | MshAT70C construction |
| VC0409-Mut-T70C-F2 | GGCATTGAATGTTTAGACTACACAGCATATAC | MshAT70C construction |
| VC0409-Mut-Seq-F1 | GGCGAAGAAAGCCAGTATTG | MshAT70C detection |

| | | |
|---|---|---|
| VC0409-Mut-Seq-R1 | CCTGCGGAGAAACTTGAATG | MshAT70C detection |
| VC2188-F1 | CCATGAGACGGTTCGTTTAC | *flaA* knockout |
| VC2188-R1-3S | CAGCGCTAATTCAGTTTAAGCGGCCATAGCGATAACGTTGTGCGGTCATC | *flaA* knockout |
| VC2188-F2-3S | GCTATGGCCGCTTAAACTGAATTAGCGCTGCAGTAGTTCACGGTACCTTC | *flaA* knockout |
| VC2188-R2 | CCAAAGATGCCGGTAAATGG | *flaA* knockout |
| VC2188-Mut-F1 | CACACTTTGGTTTCCGGTAC | FlaA mutations construction |
| VC2188-Mut-R2 | TCCGCACCATTATTGAGAGC | FlaA mutations construction |
| VC2188-Mut-A106C-R1 | TGACGCTCTGAACATGAGTTGGTACCGTTCGCCGA | FlaAA106C construction |
| VC2188-Mut-A106C-F2 | AACGGTACCAACTCATGTTCAGAGCGTCAGGCTC | FlaAA106C construction |
| VC2188-Mut-S107C-R1 | TGACGCTCACACGCTGAGTTGGTACCGTTCGCCGAT | FlaAS107C construction |
| VC2188-Mut-S107C-F2 | AACGGTACCAACTCAGCGTGTGAGCGTCAGGCTCTG | FlaAS107C construction |

| | | |
|---|---|---|
| VC2188-Mut-A106C-S107C-R1 | TGACGCTCACAACATGAGTTGGTACCGTTCGCCGAT | FlaAA106CS107C construction |
| VC2188-Mut-A106C-S107C-F2 | AACGGTACCAACTCATGTTGTGAGCGTCAGGCTCTG | FlaAA106CS107C construction |
| VC2188-Mut-E332C-R1 | CGACGCACACACGTTCTCCTGAATATTCGACAG | FlaAE332C construction |
| VC2188-Mut-E332C-F2 | ATATTCAGGAGAACGTGTGTGCGTCGAAAAGTC | FlaAE332C construction |
| VC2188-Mut-G23C-R1 | GTTAAGCTCACACGTCGCCTTGGTCAGATAACGTTGTG | FlaAG23C construction |
| VC2188-Mut-G23C-F2 | TATCTGACCAAGGCGACGTGTGAGCTTAACACCTCCA | FlaAG23C construction |
| VC2188-Mut-N26C-R1 | TCCATGGAGGTACAAAGCTCTCCCGTCGCCTTGGT | FlaAN26C construction |
| VC2188-Mut-N26C-F2 | ACGGGAGAGCTTTGTACCTCCATGGAACGCCTCTCA | FlaAN26C construction |
| VC2188-Mut-N83C-R1 | GTCGATTCACACATCGCACCTTCTGCGGTTTGAG | FlaAN83C construction |
| VC2188-Mut-N83C-F2 | AGAAGGTGCGATGTGTGAATCGACCAGCATTTTGCAGC | FlaAN83C construction |

| | | |
|---|---|---|
| VC2188-Mut-S325C-R1 | GTTCTCCTGAATATTACACAGGTTACTGATGCTGTGAC | FlaAS325C construction |
| VC2188-Mut-S325C-F2 | ATCAGTAACCTGTGTAATATTCAGGAGAACGTGGAAGCGTC | FlaAS325C construction |
| VC2188-Mut-S87C-R1 | CGCTGCAAAATACAGGTCGATTCATTCATCGCACCT | FlaAS87C construction |
| VC2188-Mut-S87C-F2 | GAATCGACCTGTATTTTGCAGCGTATGCGTGACCTC | FlaAS87C construction |
| VC2188-Mut-S376C-R1 | GTGAACTACTGCAATAAACAGATTGCAGAGTTTGGC | FlaAS376C construction |
| VC2188-Mut-S376C-F2 | TGCAATCTGTTTATTGCAGTAGTTCACGGTACCTTC | FlaAS376C construction |
| VC2188-Mut-V117C-R1 | ATCTTGCAGTGCACACGACTCTTCATTCAGAGCCTG | FlaAV117C construction |
| VC2188-Mut-V117C-F2 | GAAGAGTCGTGTGCACTGCAAGATGAACTGAACCGTA | FlaAV117C construction |
| VC2188-Mut-Seq-F1 | TGAGCTTGCGAACTCGATAG | FlaA mutations detection |
| VC2188-Mut-Seq-R1 | CGTTCTTCAGCGGATGATAG | FlaA mutations detection |
